# One size does not fit all: Family specific differences in seasonal patterns of abundance and behavior in butterfly communities

**DOI:** 10.1101/2023.05.22.541638

**Authors:** Grace E. Hirzel, Ashlyn E. Anderson, Erica L. Westerman

## Abstract

Animal communities can undergo seasonal shifts in assemblage, responding to changes in their environment. Animal behavior can also shift due to seasonal environmental variation, with the potential to shape ecosystems. However, it is unclear if similar environmental factors and time scales affect both abundance and behavior. We examined how butterfly abundance and behavior change seasonally in temperate prairies and a butterfly garden, and if the factors driving variation differ between taxonomic families. We conducted monthly abundance surveys year-round and biweekly abundance and behavior surveys during the summer and fall, in 2017-2021 and 2018-2020 respectively. We also determined how ambient light, temperature, precipitation, and time of year interact to affect butterfly abundance and behavior. We found increased temperature and light levels correlate with increases in general butterfly abundance. Unlike the greater community, Lycaenidae abundance decreased as weekly precipitation increased, and Papilionidae abundance did not respond to changes in environmental factors. Only Nymphalidae changed behavior in response to environmental factors, increasing thermoregulatory behaviors as temperature and light levels decreased. These results indicate that lineages may differ in their sensitivity to environmental factors, which could result in disproportionate changes in their abundances in response to future climate change and anthropogenic-driven disturbance.

## Introduction

Habitats can change drastically with the seasons, either to the benefit or detriment of their animal inhabitants (Campuzano et al. 2020; Wettlaufer et al. 2023). As a result, community structure can shift throughout the year, as animal populations respond to seasonal environmental changes (see Cerda et al. 1997; Desmond et al. 2002). Shifts in the community can be measured via animal abundance and diversity, revealing both intra– and inter-annual differences (Highland et al. 2016; Turley et al. 2022). Surveying animal behavior in conjunction to abundance can also provide useful information about animal communities (Toft et al. 2007; Sawyer et al. 2017). Many animals’ first responses to environmental change are alterations in behavior, which may lead to shifts in abundance, species interactions, or ecosystem services (reviewed in (Wong and Candolin 2015)). Thus, understanding changes in behavior and when they occur can help predict community health.

Seasonal shifts in community structure have been documented across many ecosystems, from terrestrial to aquatic and tropical to temperate environments. Tropical bird communities at higher elevations decrease in species richness from the breeding to nonbreeding season (Liang et al. 2021). Near shore fish communities in the north Atlantic are composed of different species in the winter and spring vs the summer and fall (Lazzari et al. 1999). Likewise in Panama, orchid bee communities also show seasonal changes in assemblages throughout the year (Roubik and Ackerman 1987). However, multiple environmental factors covary with season and may have interactive effects that are hard to detect in a laboratory setting. This can make it difficult to determine which seasonal factors are impacting community composition and abundance.

For example, abiotic factors such as temperature, light, and precipitation can co-vary and interact, affecting animal communities directly, or indirectly through biotic factors (Dai et al. 1999; Kleypas et al. 1999; Bae and Park 2019; La Sorte and Graham 2021). In marine ecosystems, temperature and pH change seasonally and can cause shifts in the dominant animal species (Dijkstra et al. 2011). Within terrestrial systems, seasonal precipitation and vegetation can affect abundance of tropical insects (Wolda 1978), and seasonal soil humidity can affect species richness in litter dwelling arthropod communities (Wiwatwitaya and Takeda 2005). Adding further complexity, response to seasonal factors may not be immediate, such as the increase in herbivorous tropical insect abundance two months after major precipitation events (Grøtan et al. 2012). Some of the environmental factors that affect abundance, such as day length or degree days, cannot be isolated from time of year in natural settings (Rohde and Pilliod 2021). However, identifying the seasonal factors that drive shifts in community assemblages is important, as many of these environmental factors may change in timing and intensity due to phenomena such as climate change and other human disturbances ((Zhang et al. 2004), reviewed in (Trenberth 2011)).

While studies of seasonal shifts in abundance are more common, animal communities also exhibit seasonal shifts in behavior. For example, large African carnivores show seasonal patterns in movement to avoid competing or dominant species (Vanak et al. 2013). In tropical environments, fruit feeding bats change their diets to take advantage of abundant food resources, becoming pollinators instead of strict frugivores (Heithaus et al. 1975). Diets amongst tropical ground dwelling frog species are also seasonal, overlapping less during the dry season compared to the wet season (Toft 1980). Animals may also change their diurnal patterns of activity throughout the year; for example, temperate water snakes are nocturnal in summer but are diurnal in cooler months (Mushinsky et al. 1980). By affecting predator-prey interactions, pollination events, and the abundances of plant and animal heterospecifics, seasonal shifts in behavior of these animal communities have the potential to shape ecosystems.

A community’s structure depends on the varied responses (including behaviors) of individual species, affecting interactions between predators, prey, and competitors (Nagelkerken and Munday 2016). Animals often change their behavior before changes in abundances occur, as is observed when animals face anthropogenic disturbance (Tuomainen and Candolin 2011). Research on behaviors of individual species highlights how understanding behavioral patterns can help predict changes to species assemblages and ecosystems. For example, bivalves are valuable ecosystem engineers and their behavior (burrowing depth) is a better predictor of larval recruitment than adult densities are ((van Gils et al. 2009; Compton et al. 2016). Predator behavior can also shape communities, as in the case of rat snakes, which are predicted to increase nest predations as nocturnal temperatures increase due to climate change (DeGregorio et al. 2015). Scaling up to examine behavioral patterns at the community level is critical to optimizing predictive models and management strategies for entire ecosystems. As environmental conditions change throughout the year, observing community behavior in a seasonal context will give us a more complete understanding of ecosystem health.

Butterflies offer a unique system in which to study both abundance and behavior at the community level. They are relatively easy to survey with minimal disturbance and will perform feeding, flying, and thermoregulating behaviors in view of human observers. The abundance and behavior of butterflies are also of special interest because of their ability and tendency to disperse long distances, carrying pollen between plant populations (Courtney et al. 1982; Ghazanfar et al. 2016). The phenology of butterfly abundance also has important implication for bird communities, as butterfly and other lepidoptera larvae make up the majority of passerine nestling diets (Kennedy 2019). Because of their ecological importance, abundance, and ease of survey, butterflies have been used as indicators of decline in biodiversity in several western European countries (Brereton et al. 2011).

Butterflies from a wide variety of habitats are sensitive to shifts in abiotic conditions and plant communities. Both yearly temperature and precipitation can explain interannual differences, with high temperatures and low precipitation causing annual declines in North American butterfly populations (Woods et al. 2008; Robinson et al. 2012). Environmental conditions can also cause butterfly abundance to fluctuate within a given year. In butterfly communities found in neotropical dry-forests and Brazilian mountain forests, seasonal changes in temperature and precipitation affect abundance (Checa et al. 2014; Beirão et al. 2021). Similarly plant diversity and number of degree days above freezing shape alpine butterfly communities (Pellissier et al. 2013). As environments shift due to changes in land use and climate, butterfly communities may also change, particularly in terms of timing of adult emergence and when peak abundance occurs.

In addition to changes in abundance, butterflies also respond behaviorally to environmental factors. For example, decreases in covarying factors light and temperature can cause corresponding decreases in courting, patrolling, and ovipositing in individual butterfly species from multiple taxonomic families (McDonald and Nijhout 2000; Ide 2002; Vives-Ingla et al. 2023). Likewise, at the community level, flying and nectaring behaviors increase in grassland butterflies at warmer temperatures (Kral-O’Brien et al. 2021). A meta-analysis of mark-recapture studies showed that time of year also affects behavior, as the time butterflies spend on reproductive behaviors changes throughout the season for a wide range of species (Vlašánek et al. 2018). These findings highlight the sensitivity of butterfly species and butterfly communities to changes in their environment. However, butterfly communities are often species rich and comprised of species from multiple taxonomic families (Simonson et al. 2001; Wittman et al. 2017; Subedi et al. 2021). For example, in temperate North America butterfly communities often consist of 25 to 50 species (Myers et al. 2012; Smith and Cherry 2014). It remains unclear how such diversity impacts both community-wide behavioral changes and family-specific responses to abiotic environmental factors and seasonal change.

A butterfly’s response to its’ environment may be associated with its taxonomic family status and broad evolutionary history. Biological differences between taxonomic families include size and wing morphology, which affect flight speed (Hawkins and Lawton 1995; Le Roy et al. 2019). Families also differ in ability to thermoregulate, with Pieridae and Papilionidae buffering against greater changes in temperature and Hesperiidae tolerating higher temperatures compared to other butterfly families (Bladon et al. 2020; Ashe-Jepson et al. 2023). Butterfly families also vary in their sensitivity to color (Stavenga and Arikawa 2006), which mediates a range of behaviors including mate choice, host plant selection, background matching, and finding nectar sources (Obara and Hidaka 1968; Snell-Rood and Papaj 2009; Pohl et al. 2011; van Bergen and Beldade 2019). Some behaviors, such as mate locating strategies, also appear to be influenced by taxonomic family (Tiple et al. 2010). Due to these family specific variations in physiology and anatomy, we predicted that we would observe seasonal patterns of abundance and behavior that differ between butterfly families.

In this long-term study we examined how both butterfly abundance and butterfly behavior change with season in Northwest Arkansas, a region that historically consisted of both Ozark woodlands and tallgrass prairies but is now being rapidly developed for urban use. In conjunction with animal surveys, we assessed how abiotic factors, including temperature, precipitation, and light environment, changed throughout the year at our prairie and botanical garden field sites. Since these factors can affect both butterflies’ ability to thermoregulate and vision mediated behaviors, we predicted that seasonal or daily changes in temperature, precipitation, or light environment would influence patterns of thermoregulation behaviors or cause lower levels of activity. Lastly, to account for other potentially important seasonal factors that we did not measure, such as day length and vegetation height and types, we asked if temperature, precipitation, or light environment interacted with time of year to correlate with changes in abundance and behavior.

## Methods

### Field Sites

Survey sites included three prairie sites and one botanical garden in Northwest Arkansas, USA. Woolsey Wet Prairie Restoration Area (Woolsey) is a 50 acre restored wet prairie owned by the city of Fayetteville, AR, located at 36°3’52’’N and 94°13’57’’W. Chesney Prairie Natural Area (Chesney) is an 80 acre tallgrass prairie under the control of the Arkansas Natural Heritage Commission located at 36°13’10’’N and 94°28’58’’W. Stump Prairie is a privately owned 20 acre remnant tallgrass prairie, located at 36°12’18’’N and 94°29’43’’ W. Woolsey is near the edge of Fayetteville surrounded by fallow fields slowly being developed into housing. Stump and Chesney are surrounded by wheat fields and pasture. The Botanical Garden of the Ozarks (BGO) is a 44 acre botanical garden with a native butterfly house in Fayetteville, AR, located at 36°08’12”N and 94°07’06”W.

### Study animals

We identified the observed adult butterflies to the lowest taxon possible, which varied depending on the species, genus, and family. Survey members varied in levels of identification expertise, but all survey members could confidently identify all animals to families Hesperiidae, Nymphalidae, Lycaenidae, Papilionidae, and Pieridae. For monthly surveys we could identify up to 38 species with confidence. (See fig. 1 for examples of butterflies from our study sites).

**Figure 1:**
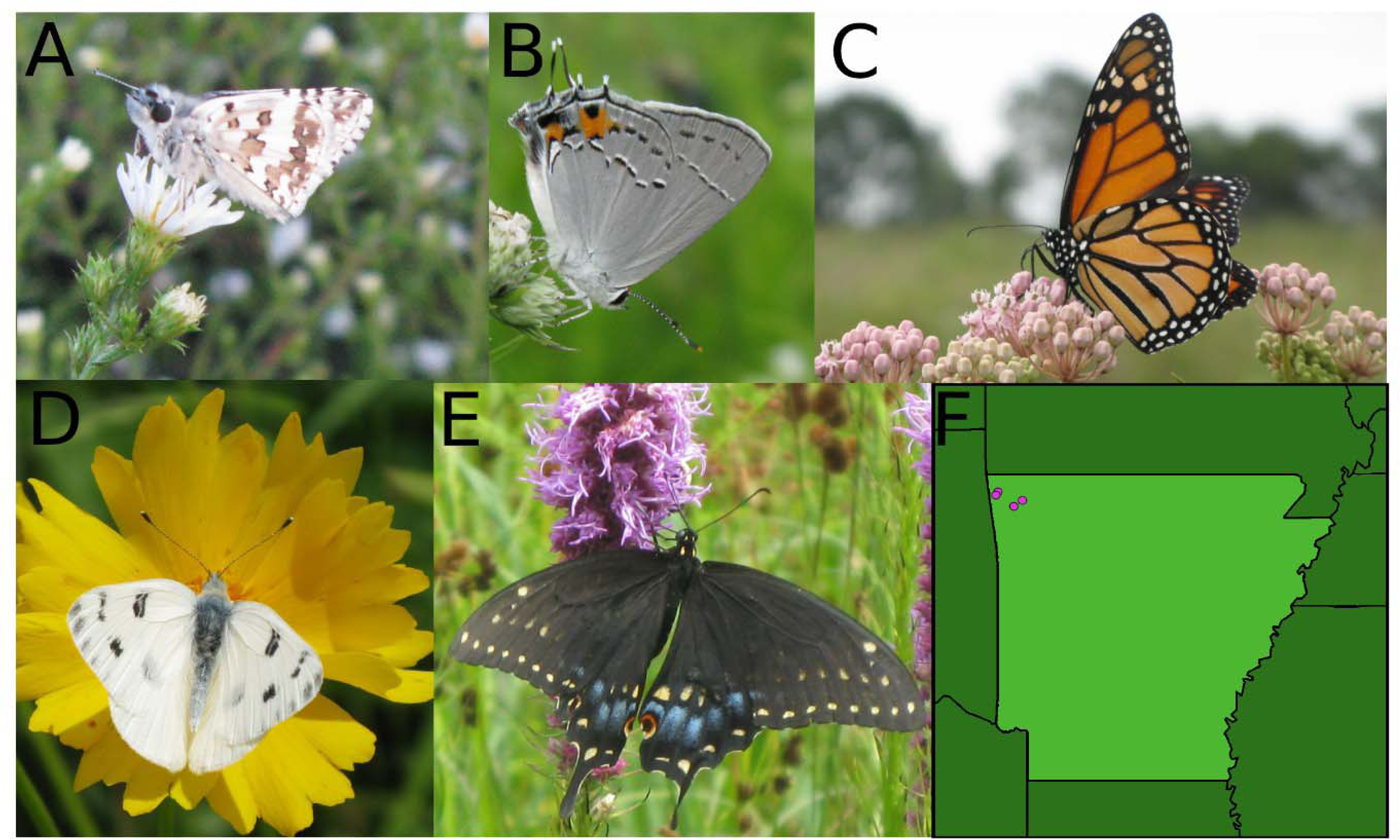
Representatives of butterfly families surveyed including Hesperiidae (A), Lycaenidae (B), Nymphalidae (C), Pieridae (D), and E) Papilionidae (E). F) Map of field sites in Arkansas.

### Transect Walks

We counted butterflies on transect surveys based on Pollard Yates Walks (Pollard & Yates 1993) to examine how the composition of butterfly communities changed throughout the season and across years. We conducted monthly surveys in small groups from July 2017 to December 2021 at the BGO and Woolsey. Survey groups usually ranged from 3 to 6 people, though in 2020 and 2021 some surveys groups were only 1 or 2 people in size because of the Covid-19 pandemic. Group surveys took place the first Thursday or Friday of the month. On days with heavy rain, we delayed surveying until the next day with suitable conditions. Surveys began at approximately 09:00 at the BGO. Surveys at Woolsey began after finishing surveys at the BGO and driving to the site, typically starting between 10:00 and 10:30. Transect walks were conducted at a slow walking pace. During these surveys we identified butterflies to the lowest taxon possible.

We also conducted surveys every other week in 2018, 2019, and 2020 from late May to early November. For these surveys, a single individual walked transects at Woolsey, Chesney, and Stump prairies. Each site had three transects. In 2018, transects were 183 meters long. In 2019 and 2020 most transects were in the same location, but lengthened by 27 meters to 210 meters. We moved Woolsey Transect 1 in 2019 after a new utility road intersected the transect. We moved Chesney Transect 3 in 2019 after analysis of 2018 data indicated that Transect 3 had significantly lower numbers of butterflies than the two other Chesney transects. We walked transects in the span of approximately 10 minutes, but times varied between 8 and 10 minutes, depending on number of butterflies. For biweekly surveys we conducted surveys in the time period between solar noon and 150 minutes after solar noon. During a survey week we visited every site once. We counted butterflies observed within 5 meters to the front and sides of the observer during biweekly transects.

### Behavior

We collected behavioral data to investigate how seasonal and other environmental conditions affected the behavior of different butterfly taxa. We recorded behavior during our biweekly visits to Woolsey, Chesney, and Stump prairies in 2019 and 2020. During the first transect we walked for each visit, we recorded the first behavior observed for every butterfly we counted.

We categorized behavior as *flying*, *resting*, *basking*, *nectaring*, *courting*, *chasing*, *rejecting*, *mating*, *hovering*, *ovipositing*, and *startled*. Butterflies were categorized as *chasing* whether they were following other butterflies or being followed. While *chasing* behavior can be part of courting in some species, for this study we defined *courting* as butterflies hovering and fluttering over another butterfly (Scott 1975). *Rejecting* was defined as butterflies that were fluttering their wings while being courted (Scott 1975). We recorded *resting* behavior when we observed inactive butterflies that had closed wings and *basking* behavior when we observed inactive butterflies that had open wings. We recorded butterflies as *startled* when we observed butterflies flying from a stationary position or flying within 0.5 meters from the observer in thick vegetation and could have been startled from a stationary position.

### Abiotic factors

To examine how seasonal light environment affected abundance and behavior of butterflies from different families, we measured absolute solar irradiance during visits to sites with a Jaz spectrophotometer (Ocean Optics). During monthly visits we measured irradiance at four predetermined spots along our transects at both the BGO and Woolsey that stayed consistent across all years. For biweekly surveys, we included an additional spot for measurements at Woolsey and also took irradiance measurements at five predetermined locations along our transects at Chesney and Stump prairies. Locations for irradiance measurements mainly stayed consistent throughout the study; however, we moved one measurement location at Chesney between 2018 and 2019. Instead of measuring irradiance at the beginning of the Transect 3 we walked in 2018, we took measurements at the end of the new Transect 3 walked in 2019 and 2020. At each designated irradiance location, we recorded three irradiance measurements. (We took up to six measurements if we suspected technical problems with saving measurements.)

We also examined how temperature and precipitation affected butterfly abundance and behavior. We used temperature and precipitation data captured by a regional weather station at Northwest Arkansas National Airport that is available online through NOAA (https://www.ncei.noaa.gov/cdo-web/datatools/lcd). For monthly surveys we used the temperature measured immediately before conducting the surveys (around 08:50 for BGO and 09:50 for Woolsey). For biweekly surveys we used the temperature measurements around 13:50, because this time always fell within the start and end points of a survey visit. For precipitation we used total levels of precipitation that fell for the Julian Week of the survey.

### Data Processing and Statistical Analysis

Irradiance data was extracted and processed using R 3.6.3 (R Core Team 2018). Using the *photobiology* and *photobiologyInOut* packages (Aphalo 2015), we extracted properties of ambient light (UV and total solar irradiance levels). These packages extracted the calibrated (“Processed”) data in the rightmost column of the *.JazIrrad files. After extracting data we used the e_irrad() function to find irradiance values for all wavelengths from 290 to 800 nm and 290 to 400 for total irradiance and ultraviolet wavelengths (UV), respectively. We determined relative levels of UV by dividing the UV irradiance by total irradiance. We averaged the measurements taken at a single measurement location during each survey visit. When comparing irradiance values to abundance and behavior, all site measurements were then averaged together.

We completed subsequent analysis of abundance and environmental data with R 4.2.2 (R Core Team 2018). Before we tested whether environmental factors, including time of year, total solar irradiance, relative UV levels, temperature, weekly precipitation, survey year, and site, affected abundance or behavior, we ran a correlation matrix to determine the correlations between these environmental factors. Correlated factors (R^2^ > 0.20) were then used in a principal component analysis. Sampling dates with missing data (eg. weather or abundance) were dropped from the analysis. Principal Components (PCs) were included in generalized linear models with the glm() function from the *lme4* package (Bates et al. 2015) to determine the effects of our PCs on abundance (monthly and biweekly data) and behavior (biweekly) of the entire butterfly community, as well as the abundance and behavior of butterfly members of each of the five families we observed (Hesperiidae, Nymphalidae, Lycaenidae, Papilionidae, and Pieridae). Models were run with a quassipoisson distribution and chi square test. To account for multiple testing we used a Bonferroni correction (α = .05, *p* = 0.0016). To further examine behavioral responses to specific factors and to visualize behavioral data we ran posthoc nominal regressions in JMP Pro 17.0.0 (JMP 2021). Non behavioral data were visualized with *ggplot2* in R (Wickham 2016).

### Ethical Statement

No butterflies were handled during this research. Permission to survey at Stump was granted by Ozark Ecological Restoration Inc and permission to survey at Woolsey was provided by Environmental Consulting Operations, Inc. We surveyed at Chesney with permission from the Arkansas Natural Heritage Commission under research permits S-NHCC-18-023, S-NHCC-19-015, and S-NHCC-20-010. The BGO permitted us to survey on their property.

## Results

We walked 109 monthly survey transects from 2017-2021 and saw species from 5 butterfly families (see fig. 1). We counted 2,919 Nymphalidae, 1,141 Hesperiidae, 837 Lycaenidae, 606 Pieridae, and 262 Papilionidae (fig. S1A) We walked 342 biweekly survey transects from 2018-2020. We counted a total of 3,559 Nymphalidae, 1,195 Hesperiidae, 561 Lycaenidae, 373 Pieridae, and 95 Papilionidae (fig. S1B).

### Abiotic factors covary and change throughout the year

Many, but not all, of the abiotic factors we measured changed throughout the year. Ambient temperature was highest from June to September in monthly surveys (fig. 2A) and highest in July for biweekly surveys (fig. 2B). Precipitation during the week of the survey did not change significantly from month to month or week to week for either monthly or biweekly surveys (fig. 2C, D). Total irradiance was highest in June and August in monthly surveys and highest for the entire period from May through July in biweekly surveys (fig. 2E, F). Relative UV did not change significantly throughout the year for monthly surveys, but changed seasonally for biweekly surveys, with relative UV levels peaking at the end of May (fig 2G, H).

**Figure 2:**
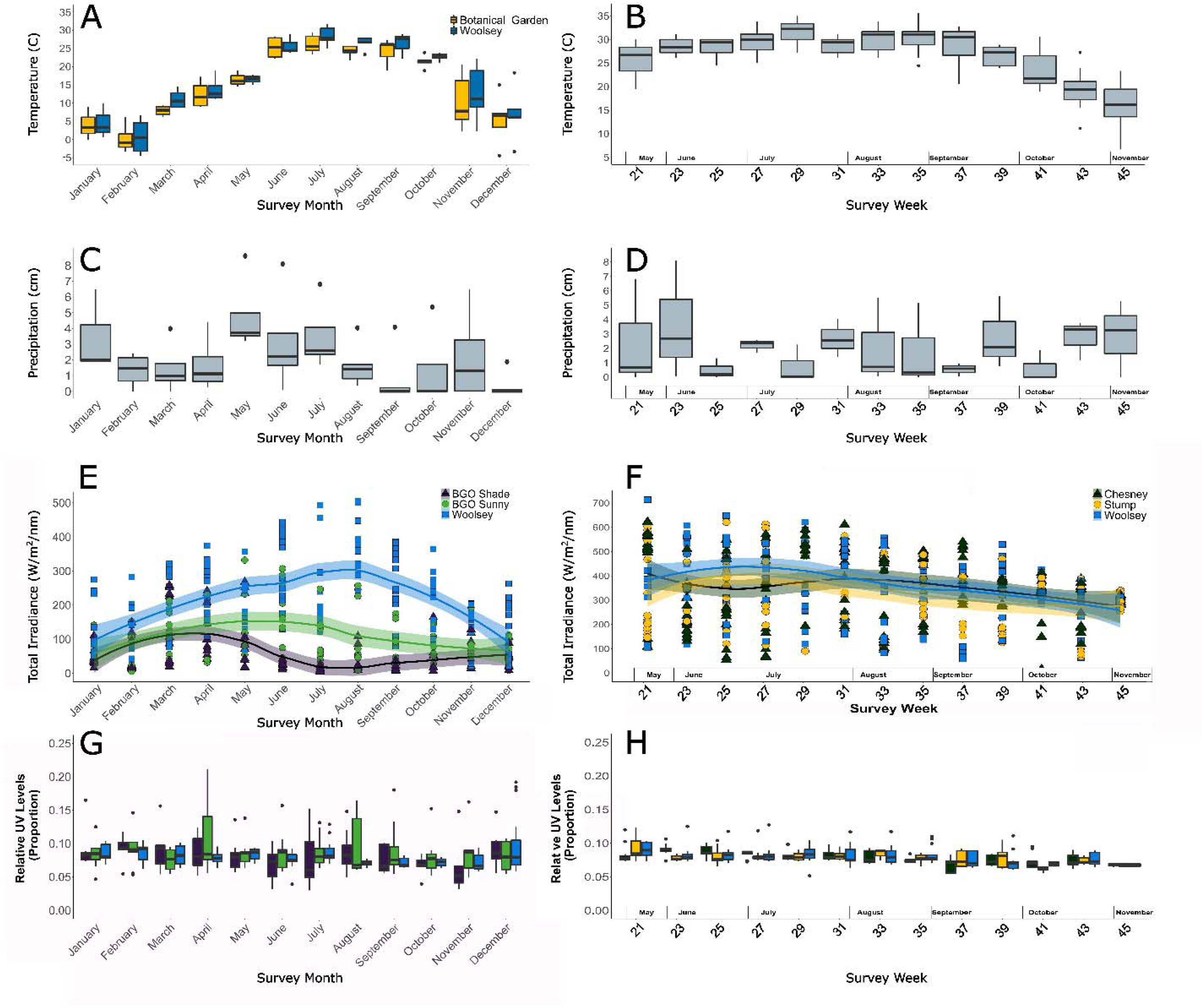
Abiotic factors for monthly and biweekly surveys. Temperatures from a local regional airport from the hour closest to survey time for both monthly (A) and biweekly (B) surveys. Precipitation levels from a local regional airport from the week proceeding survey weeks for both monthly (C) and biweekly (D) surveys. Absolute irradiance measurements for monthly (E) and biweekly (F) surveys and relative UV measurements for monthly (G) and biweekly (H) surveys that were collected at field sites.

Time of year and weather conditions can be highly correlated. To investigate how these factors interacted and affected butterfly abundance and behavior, we ran principal component analyses. Survey time of year (month/week), survey year, temperature, precipitation, total irradiance, and relative UV were correlated with each other for both monthly (table S1) and biweekly (table S2) surveys and were included in principal component analyses. Site was correlated to total irradiance in monthly surveys, but we did not include it in the principal component analysis. Our two sites, Woolsey and the BGO, differ in factors other than light environment, such as plant species and human disturbance. To better understand differences in abundance between sites, we instead included site as an independent variable in our models. In biweekly surveys, site did not correlate to any of the other variables we measured and was included as an independent variable in our models.

### Butterfly abundance increases with absolute irradiance and temperature in year round surveys

Principal Component 1 (PC1), PC2, and PC3 accounted for 75.3% of the total variance in abiotic data for monthly surveys (table 1). PC1 accounted for 32.2% of variance and primarily comprised of equal loadings of temperature, total irradiance, and negative relative UV. Lower weightings were given to year and survey months, with little impact of precipitation. We interpreted this to mean that high PC1 values represent “sunny conditions.” PC2 accounted for 23.7% of variance and predominately comprised of equal loadings of survey month and survey year, which were negatively correlated. Lower weightings were given to temperature, total irradiance, and precipitation, with little impact of relative UV. PC3 accounted for 19.5% of variance and comprised largely of increasing weekly precipitation. Other factors were equally loaded and contributed to a lesser extent.

**Table 1:** Loadings for principal components analysis of monthly survey abiotic data

|  | PC1 | PC2 | PC3 | PC4 | PC5 | PC6 |
| --- | --- | --- | --- | --- | --- | --- |
| Year | 0.314 | -1.616 | -0.284 | 0.405 | 1.400 | 0.284 |
| Survey Month | 0.289 | 1.456 | -0.341 | 0.656 | -2.459 | 0.154 |
| Temperature | 0.598 | 0.824 | 0.361 | 0.250 | 4.310 | -0.038 |
| Weekly Precipitation | -0.134 | -0.504 | 0.805 | 0.633 | -2.085 | 0.032 |
| Total Irradiance | 0.631 | 0.085 | 0.301 | -0.783 | -2.031 | 0.218 |
| Relative UV | -0.699 | 0.755 | 0.158 | -0.161 | 1.868 | 0.350 |
| Proportion of Variance | 0.322 | 0.237 | 0.195 | 0.125 | 0.069 | 0.054 |
| Cumulative Proportion | 0.322 | 0.558 | 0.753 | 0.877 | 0.946 | 1.000 |

For models of monthly survey abundance, we included three principal components (PC1, PC2, and PC3) and site. PC1 had a positive effect on total butterfly abundance (fig. 3A, table 2). When data were analyzed by Lepidoptera family, we recovered a strong positive correlation between PC1 and Hesperiidae, Nymphalidae, and Pieridae abundances, with other families showing significant but weak (<0.1) correlations (fig. 3D, 3G, 3J, 3M, 3P, table 2). PC2 significantly affected butterfly abundance for most families, but changes in PC2 did not strongly correlate to changes in abundance (R^2^ less than 0.1 for all families) (fig. 3B, 3E, 3H, 3K, 3N, 3Q, table 2). PC3 (precipitation) had a negative effect on Lycaenidae abundance and had significant weak (<0.1) effects on Nymphalidae and Pieridae (fig. 3C, 3F, 3I, 3L, 3O, 3R, table 2). Butterfly abundance was significantly higher at Woolsey than at the BGO for families Hesperiidae and Papilionidae (fig 4A, table 2).

**Figure 3:**
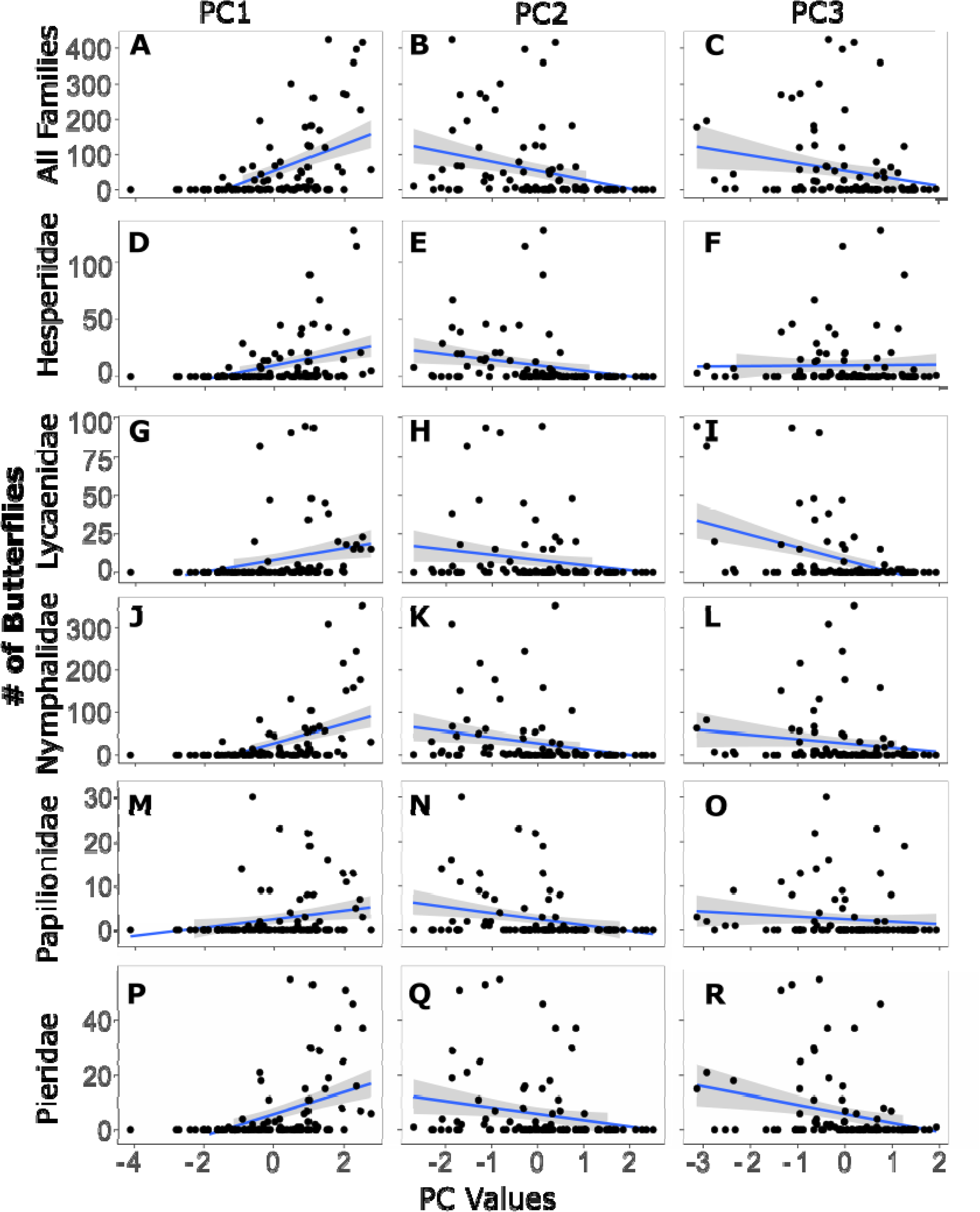
Effect of abiotic factors on butterfly abundance. A) Overall butterfly abundance increases as PC1 (temperature and absolute irradiance vs relative UV levels) increases, but does not strongly correlate with changes in PC2 (B) or PC 3 (C). Hesperiidae, Nymphalidae, and Pieridae abundance increase with PC1 (D, J, P), but do not strongly correlate with PC2 (E, K, Q) or PC3 (F, L, E). Lycaenidae abundance does not correlate strongly with PC1 (G) or PC2 (H), but decreases as PC3 (weekly precipitation) increases (I). Papilionidae abundance did not correlate strongly with any PC (M-O).

**Figure 4:**
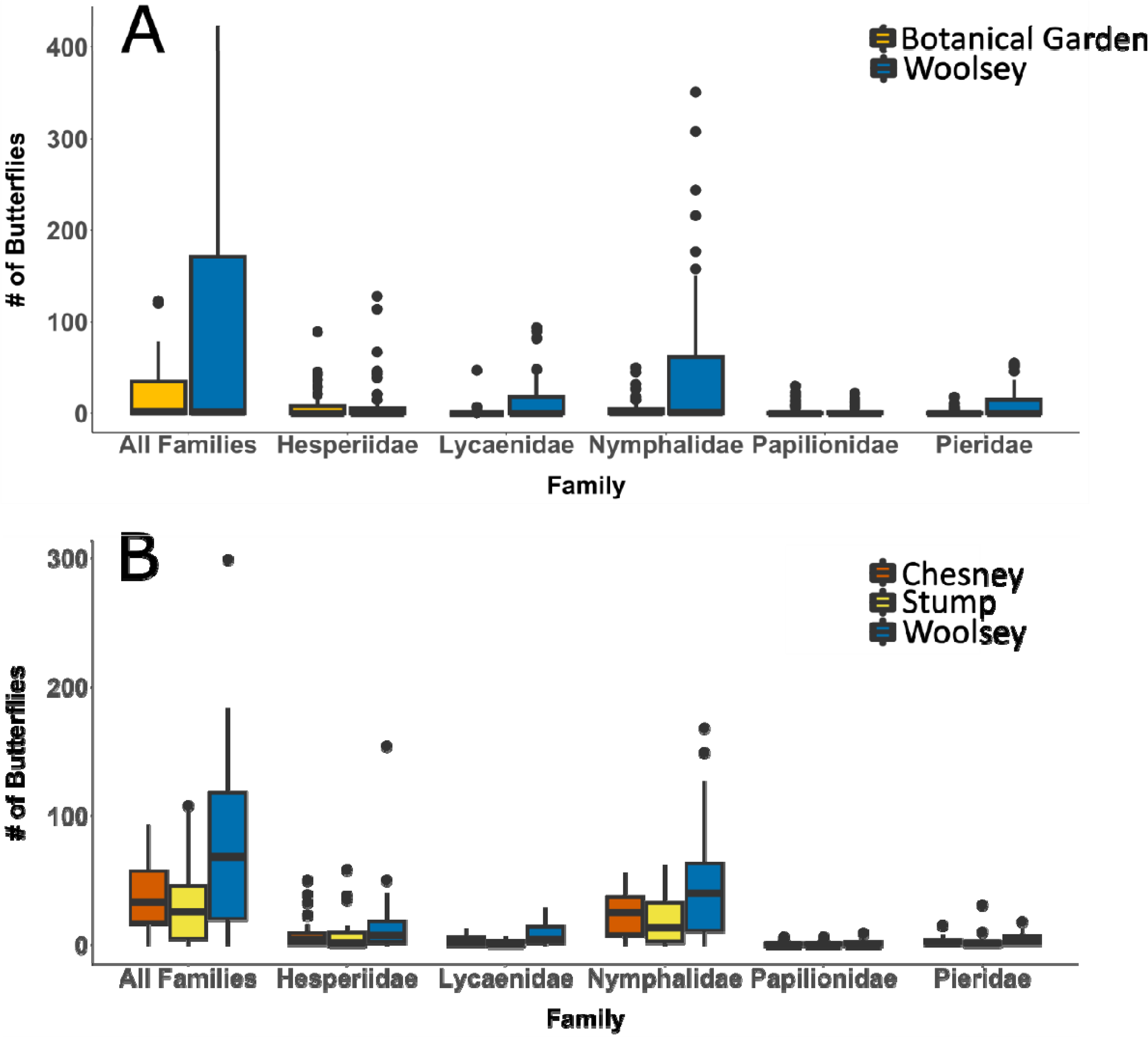
Butterfly abundance by site. Woolsey had more butterflies in both monthly (A) and biweekly (B) surveys.

**Table 2:**
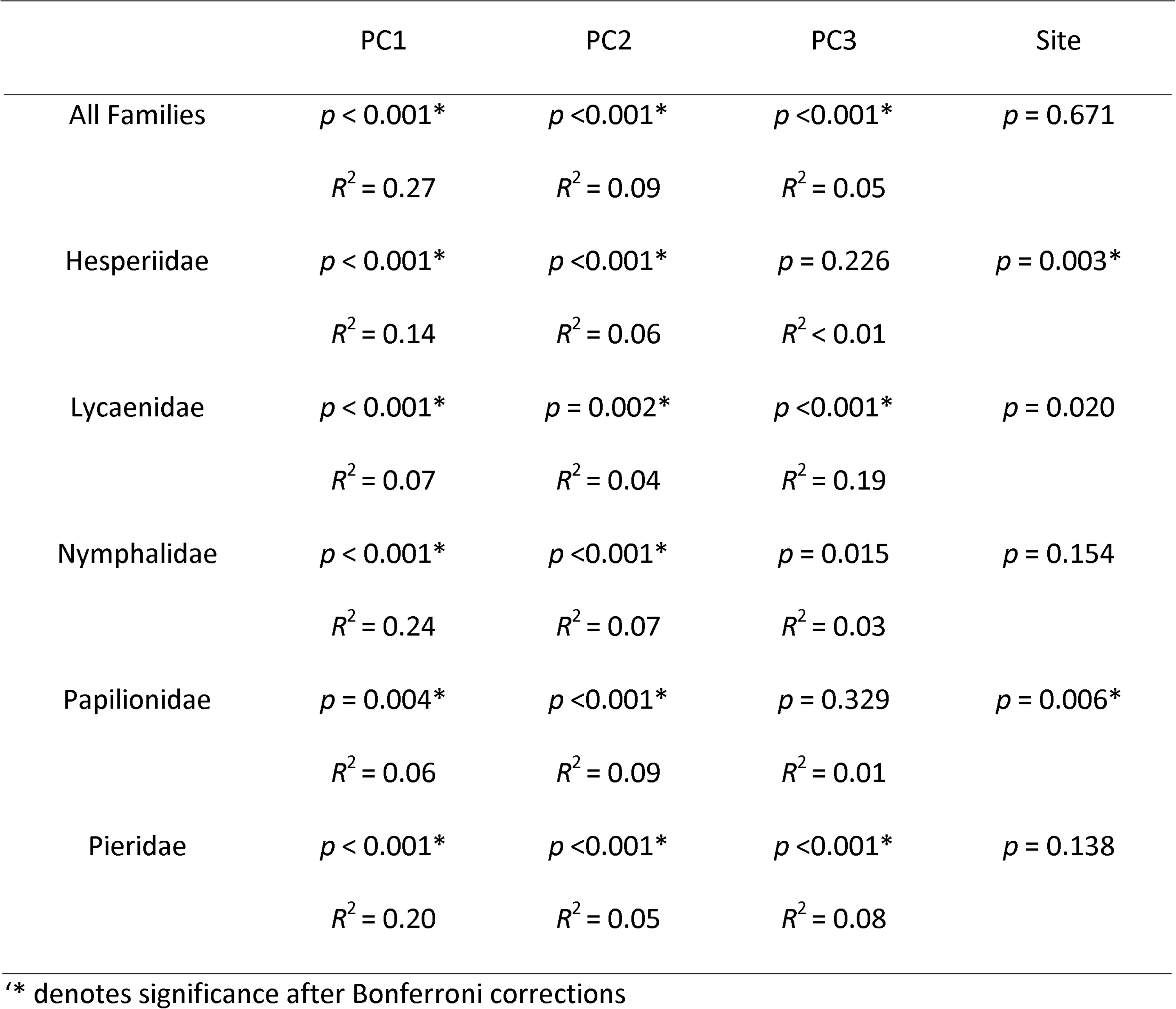
Generalized linear model analyses of monthly survey abundance data

### Time of year, temperature, and light cause changes in butterfly abundance in surveys conducted during the flight period

In our biweekly surveys, four principal components explained 89.9% of variance of abiotic data (table 3). Principal Component 1 (PC1) accounted for 32.7% of variance and was comprised of equal loadings of temperature, total irradiance, relative UV, and negative survey week. We interpreted PC1 to represent “seasonality.” PC2 accounted for 28.0% of variance and comprised primarily of weekly precipitation, relative UV, and negative total irradiance and survey year. We interpreted PC2 to represent “cloud and rain levels.” PC3 accounted for 15.0% of variance and was comprised primarily of survey year. PC4 accounted for 14.2% of variance and comprised of primarily weekly precipitation.

**Table 3:** Loadings for principal components analysis of biweekly survey abiotic data

|  | PC1 | PC2 | PC3 | PC4 | PC5 | PC6 |
| --- | --- | --- | --- | --- | --- | --- |
| Year | -0.468 | 1.513 | 1.124 | 0.004 | -0.125 | 0.139 |
| Survey Week | -1.570 | 0.446 | -0.355 | 0.030 | 0.496 | 0.319 |
| Temperature | 1.343 | 1.219 | 0.135 | 0.007 | 0.831 | -0.036 |
| Weekly Precipitation | -0.196 | -1.752 | 0.287 | 0.737 | 0.192 | -0.054 |
| Total Irradiance | 0.906 | 1.889 | -0.391 | 0.423 | -0.386 | 0.252 |
| Relative UV | 0.986 | -2.316 | 0.201 | -0.201 | -0.008 | 0.380 |
| Proportion of<br>Variance | 0.327 | 0.280 | 0.150 | 0.142 | 0.075 | 0.026 |
| Cumulative<br>Proportion | 0.327 | 0.607 | 0.758 | 0.899 | 0.974 | 1.000 |

We included five factors in our models of biweekly abundance (PC1, PC2, PC3, PC4, and site). Only PC1 significantly affected and had a strong correlation with butterfly abundance. As PC1 increased, total butterfly abundance decreased (fig. 5A, table 4). This was also observed for families Lycaenidae and Nymphalidae when families were analyzed individually (fig. 5B-F, table 4).

**Figure 5:**
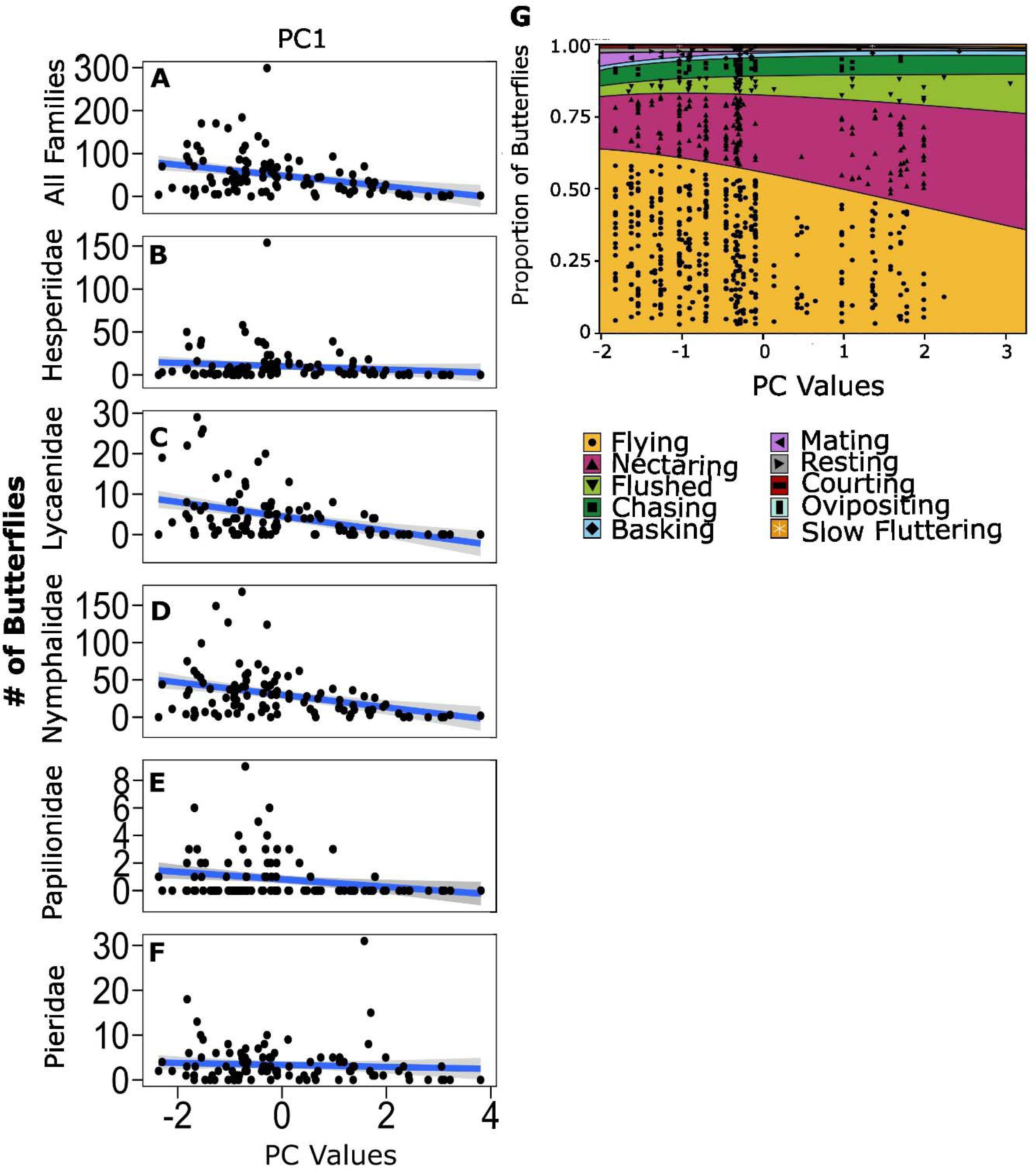
Effect of PC1 (temperature vs survey week) on biweekly butterfly abundance but not behavior. As PC1 decreases in biweekly surveys butterfly abundance decreases overall (A) and for families Lycanidae (C) and Nymphalidae (D), but not for families Hesperiidae (B), Papilionidae (E), and Pieridae (F). G) PC1 did not affect behavior.

**Table 4:** Generalized linear model analyses of butterfly abundance during biweekly surveys

|  | PC1 | PC2 | PC3 | PC4 | Site |
| --- | --- | --- | --- | --- | --- |
| All Families | $p < 0.001^*$<br>$R^2 = 0.13$ | $p = 0.013$<br>$R^2 = 0.03$ | $p = 0.311$<br>$R^2 < 0.01$ | $p = 0.755$<br>$R^2 < 0.01$ | $p < 0.001^*$ |
| Hesperiidae | $p = 0.066$<br>$R^2 = 0.02$ | $p = 0.013$<br>$R^2 = 0.03$ | $p = 0.561$<br>$R^2 < 0.01$ | $p = 0.265$<br>$R^2 < 0.01$ | $p = 0.107$ |
| Lycaenidae | $p < 0.001^*$<br>$R^2 = 0.17$ | $p = 0.348$<br>$R^2 < 0.01$ | $p = 0.754$<br>$R^2 < 0.01$ | $p = 0.009$<br>$R^2 = 0.02$ | $p < 0.001^*$ |
| Nymphalidae | $p < 0.001^*$<br>$R^2 = 0.14$ | $p = 0.094$<br>$R^2 = 0.01$ | $p = 0.295$<br>$R^2 < 0.01$ | $p = 0.821$<br>$R^2 < 0.01$ | $p < 0.001^*$ |
| Papilionidae | $p = 0.003^*$<br>$R^2 = 0.06$ | $p = 0.078$<br>$R^2 = 0.02$ | $p = 0.490$<br>$R^2 < 0.01$ | $p = 0.918$<br>$R^2 < 0.01$ | $p = 0.483$ |
| Pieridae | $p = 0.439$<br>$R^2 < 0.01$ | $p = 0.011$<br>$R^2 = 0.06$ | $p = 0.527$<br>$R^2 < 0.01$ | $p = 0.571$<br>$R^2 < 0.01$ | $p = 0.200$ |

\* denotes significance after Bonferroni corrections.

### Basking behavior changes with light and temperature, but only in Nymphalidae

We included these same five variables (PC1, PC2, PC3, PC4, and Site) in our models when analyzing our biweekly behavioral data. When analyzing the butterfly community as a whole, only PC2 affected behavior (table 5). At lower levels of PC2, butterflies were more likely to be seen basking; at higher levels they were more likely to be seen nectaring (fig. 6G). When families were analyzed individually, PC2 also affected Nymphalidae behavior, but not the behavior of the other families (table 5). Nymphalidae behavior mirrored the overall community trend and were more likely to be seen basking at lower PC2 levels and more likely to be seen nectaring at higher PC2 levels (fig. 6H). None of the principal components significantly affected Hesperiidae or Lycaenidae behavior (table 5). There were not enough data to analyze the behavior of Pieridae or Papilionidae individually.

**Figure 6:**
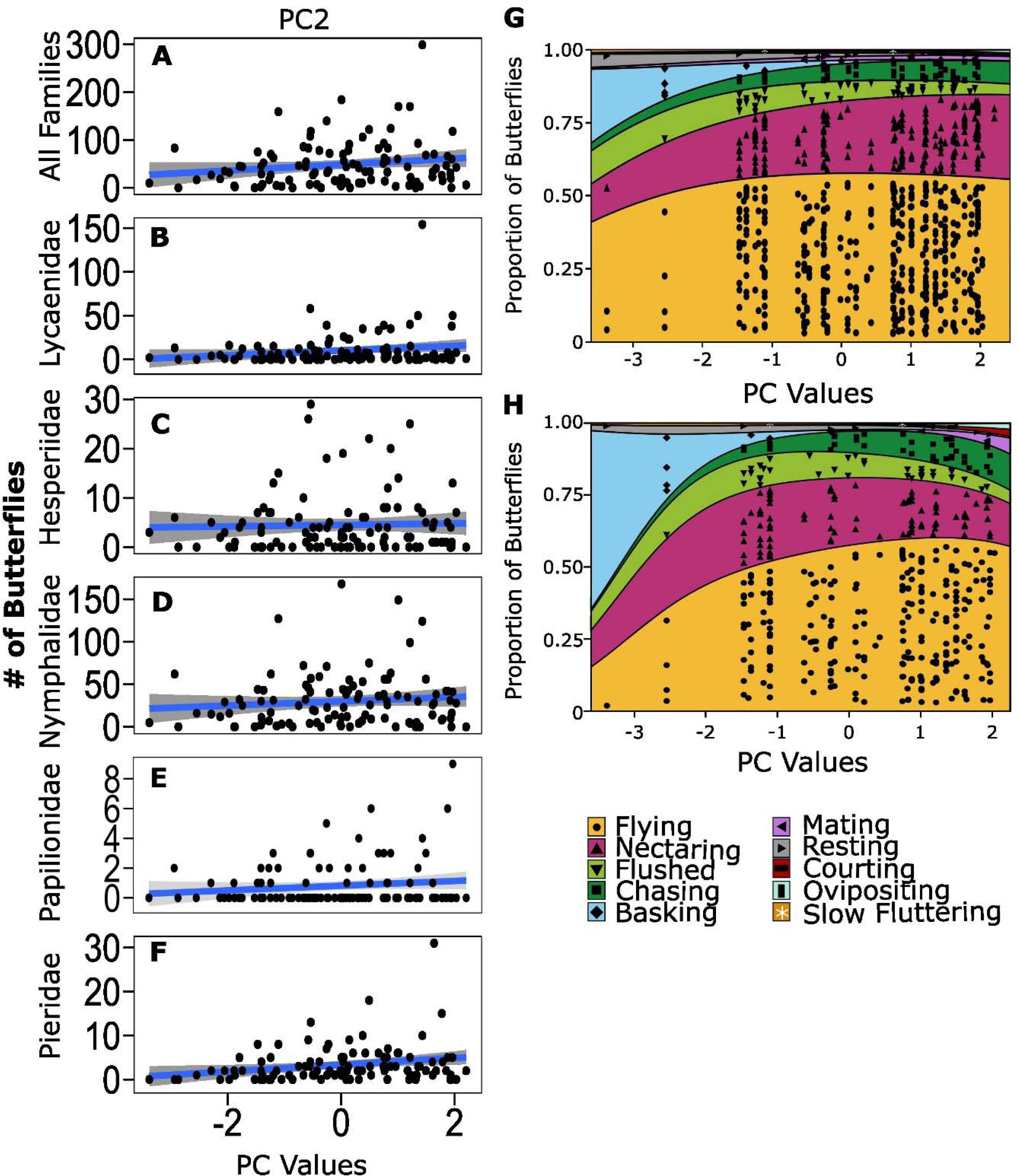
Effect of PC2 (year/temperature and absolute irradiance vs weekly precipitation and relative UV levels) on biweekly butterfly behavior but not abundance. Changes in biweekly butterfly abundance do not correlate with changes in PC2 (A-F). However, as PC2 increases basking behavior decreases overall (G) and in Nymphalidae (H).

**Table 5:** Generalized linear model analyses of behavior during biweekly surveys

|  | PC1 | PC2 | PC3 | PC4 | Site |
| --- | --- | --- | --- | --- | --- |
| All Families | $p = 0.822$ | $P < 0.001^*$ | $p = 0.122$ | $p = 0.037$ | $p = 0.344$ |
| Hesperiidae | $p = 0.020$ | $p = 0.683$ | $p = 0.560$ | $p = 0.561$ | $p = 0.368$ |
| Lycaenidae | $p = 0.468$ | $p = 0.358$ | $p = 0.374$ | $p = 0.568$ | $p = 0.150$ |
| Nymphalidae | $p = 0.139$ | $p < 0.001^*$ | $p = 0.177$ | $p = 0.851$ | $p = 0.245$ |

\* denotes significance after Bonferroni corrections.

## Discussion

We found that seasonal environmental factors affect butterfly abundance and behavior in wet and dry prairie communities as well as in botanical gardens. In the course scale (monthly) year-long surveys, we found that as temperature and sunny conditions increase, total butterfly abundance increases. Both these abiotic factors affected abundance more than time of year in the monthly surveys. We observed a similiar pattern in fine-scale (biweekly) sampling efforts during the flight period (late spring through fall); biweekly abundance decreased in response to the covarying decrease in temperature and total irradiance as the flight period progressed. Community-wide patterns were reflected in Nymphalidae. Hesperiidae, Lycaenidae, and Pieridae abundances also followed community wide patterns but only in one of the survey data sets. Lycaenidae abundance was also negatively affected by increased precipitation. Conversely, Papilionidae did not respond to any of the environmental conditions we measured.

We also found seasonal effects on community-wide butterfly behavior, with butterflies as a whole decreasing basking behavior as temperature and light increase and relative UV levels decrease. However, when we examined families individually, we found that only nymphalids show changes in behavior in response to any of the abiotic factors we measured. When combined, our results suggest that while temperature and light affect both overall butterfly abundance and behavior, butterfly family (and associated evolutionary history) may play an important role in how butterflies respond to their environments.

Our finding that butterfly communities in mid-latitude prairies and botanical gardens change in abundance with changes in temperature may reflect global patterns and complexities encountered when assessing communities in other habitats and seasonal temperature regimes. In both tropical and temperate ecosystems butterfly abundance generally increases as temperatures increase (Checa et al. 2009; Rohde and Pilliod 2021). We saw this reflected in our analyses of monthly data as abundance increases with temperature. In our biweekly surveys, time of year also is important, since butterfly abundance decreased as the season progressed and temperature decreased. Time of year and temperature interact to affect abundance in other butterfly communities. In tropical systems, butterflies respond to interactions between temperature and seasonal precipitation, with abundance increasing or decreasing with precipitation levels depending on the habitat type (Checa et al. 2009; Beirão et al. 2021). Interactions between time of year and temperature appear in other North American grasslands, where day length and growing degree days interact with temperature to not only affect abundance of butterflies and moths, but also abundance of bees and flies (Rohde and Pilliod 2021). Interactions between temperature and other seasonal factors are not limited to butterfly and terrestrial communities: in marine ecosystems temperature interacts with pH to cause seasonal changes in invertebrate communities (Dijkstra et al. 2011). In our study survey month and survey week served as proxies for important seasonal factors that we might not have considered. Although we measured seasonal light and weekly precipitation, animal communities can also be affected by such seasonal factors as vegetation and soil moisture (Wolda 1978, Wiwatwitaya and Takeda 2005). Future studies could examine whether seasonal temperature interacts with other aspects of seasonality and time of year to differentially affect certain groups of ectotherms and other temperature sensitive animals within their communities.

We also found that butterfly abundance increases with total light and decreases with relative UV. Changes in solar irradiation and canopy cover can explain changes in butterfly abundance across different habitats and within forest habitats; whether the effect of light is negative or positive depends on the scale and habitat type (Waltz and Covington 2004; Houlihan et al. 2013; Grundel et al. 2020). Here we show that light environment also covaries with time of year and other aspects of seasonality to affect butterfly abundance, and that these effects differ for relative UV light and total irradiance. In the monthly surveys increased relative UV could have been from either increased shade or cloud cover, since some surveys took place at a botanical garden with shaded areas. In our biweekly surveys increased relative UV was only from increased cloud cover, as these prairie sites lacked shade. However, increased relative UV across all sites caused a decrease in butterfly abundance, indicating that regardless of cause, butterflies respond to this aspect of light environment. Previous studies show butterfly abundance changes with qualitative or estimated shifts in canopy and cloud cover (Fischer and Fiedler 2001; Grundel et al. 2020). By measuring quantitatively, we were able to show that canopied and cloudy environments may affect animals similarly, possibly due to changes in their visual environment. Changes in canopy cover also influence changes in abundance in other animals capable of seeing UV, including birds and other insect pollinators like bees and flies (McCabe et al. 2019) (Patten and Smith-Patten 2012). Often changes in canopy cover are framed in the context of changing temperature, it would be interesting to examine how communities of other animals respond to relative UV, which we show is an important aspect of increased shade and cloud cover. This is especially important now, since canopy cover is decreasing in the tropics and increasing in other parts of the globe, causing wide-ranging shifts in light environments (Song et al. 2018).

While temperature and light environment affected the butterfly community at large, we found it particularly interesting that they did not affect the abundances of all families equally and had no effect on the abundance of Papilionidae in either the monthly or biweekly surveys. In general, Papilionidae are better at thermal buffering (maintaining specific body temperature) than other butterfly species (Ashe-Jepson et al. 2023). Thermal buffering may be useful when temperatures fluctuate, but is also associated with a narrower thermal niche, meaning that higher temperatures may affect Papilionidae more negatively than other families (Ashe-Jepson et al. 2023). This may indicate that climate change will affect populations of species in Papilionidae in different ways than populations of species in other butterfly families. In general, butterflies have a relatively narrow thermal niche compared to other, smaller insect pollinators which may thrive at cooler temperatures that butterflies cannot tolerate (Kühsel and Blüthgen 2015). However, the differences in family responses to temperature may be more indirect; for example, bird species that eat animals decline in abundance with colder winter temperatures while other bird species do not (Reif et al. 2010). Butterflies also have a wide variety of diets, both as larvae and adults (Dyer et al. 2007, Krenn 2008, Opler and Krizek 1984). If host plant abundance changes due to environmental change, butterfly populations may respond in kind (Yamamoto et al. 2007). Future studies could explore the mechanisms behind family specific responses to temperature and light, whether it be physiological or indirectly caused, for example by diet.

Lycaenidae also expresses a family specific response to abiotic factors, as it is the only family in the monthly surveys that decreases in abundance as weekly precipitation increases. In studies taking place in the tropics, butterfly abundance increases due to precipitation events, but this increase can lag behind precipitation events by two months (Grøtan et al. 2012). Yearly precipitation can also increase overall butterfly abundance in temperate habitats (Robinson et al. 2012). In contrast to these studies, our analysis examined the more immediate response of butterflies to precipitation. Lycaenidae contains some of the smallest butterflies in our area, so this family may be more sensitive to certain conditions than other families containing larger butterflies (Spencer 2006, Bladon et al. 2020, Ashe-Jepson et al. 2023). It will be important to determine if Lycaenidae are more greatly impacted by precipitation events, especially since precipitation is predicted to increase at higher latitudes and decrease at lower latitudes in current climate models (Trenberth 2011). Further research can explore the possible physiological mechanisms that are disproportionally affecting Lycaenidae compared to larger butterflies or those from other families.

Butterfly behavior also responded to environmental conditions. Unlike what we observed for biweekly abundance, biweekly behavior was more strongly affected by temperature and light, rather than time of year. Still, biweekly behavior and monthly abundance both were affected by similar conditions. Behavioral surveys could be used in future surveys as indicators of community health, if abundance and behavior respond to similar factors. Other animals, including other pollinators, change their behaviors seasonally (Vanak et al. 2013, Heithaus et al. 1975, Toft 1980). We found a seasonal change in *basking*, a thermoregulatory behavior. Like in our butterflies, seasonal behaviors in other ectotherms may be related the thermoregulation (Mushinsky et al. 1980). In general, while behavioral changes may result in changes to abundance of the study taxon (van Gil et al. 2009, Tuomainen & Candolin 2011), there may be greater implications for other species in the ecosystem, especially if the study taxon are important predators, prey, or competitors (DeGregorio et al. 2015, Reviewed in Nagelkerken & Munday 2015). Surveying seasonal behavior may help describe mechanisms behind behavioral change and in the future, predict changes in abundance.

The butterfly community at large responded behaviorally to changes in environmental conditions, though this was only reflected in Nymphalidae at the family level. Nymphalid behavior changed significantly in response to environmental factors: mainly temperature, weekly precipitation, total irradiance and relative UV. In contrast to nymphalid abundance, nymphalid behavior was not strongly affected by time of year. In our area Nymphalidae is the most speciose family (Spencer and Simons 2006), so it could be that patterns of nymphalid behavior are due to covarying shifts in nymphalid assemblages. An alternate hypothesis is that the community changes its behavior in response to different environmental conditions, while remaining relatively stable in composition. Many Nymphalidae species are sensitive to changes in environmental conditions and exhibit seasonal wing colors and behaviors (Brakefield and Reitsma 1991; Smith 1991; Davis 2009; McElderry 2016; Järvi et al. 2019). We suspect the observed Nymphalidae-wide changes in behavior are probably due to both proposed mechanisms: intraspecific plasticity and changes in community assemblage. However, since different principal components are responsible for changes in behavior and abundance and only abundance was affected by survey site, it is unlikely that changes in assemblage alone cause changes in behavior. Future studies could target specific species to compare between families and determine mechanisms. If butterfly families differ in response to light and temperature conditions, this has important implications for butterfly conservation and management strategies, especially if family specific differences are due to differing sensitivity to environmental factors.

## Conclusions

We found that seasonal conditions affect both butterfly abundance and behavior. This indicates that the degree to which communities are affected by seasonal change is not just a matter of how many animals are present, but also what they are doing. Scale of sampling affected our results: while temperature and light were better predictors of abundance for the entire year during monthly sampling, this was not true when we surveyed animals biweekly during the time of year that they were most abundant. Instead, survey week (a proxy for unmeasured seasonal conditions) and temperature were better predictors of biweekly butterfly abundance. Our data suggest that some butterfly families are more sensitive to ambient environmental conditions than other others, a hypothesis that could be tested in future comparative studies. Examining how seasonal conditions influence family specific responses in behavior and abundance may help predict which groups of animals are most vulnerable as climate change and habitat degradation cause abiotic factors to shift.

## Acknowledgements

We thank all who participated in monthly surveys over the years of this project: Dylan Meyer, Deonna Robertson, Timothy Sullivan, Peyton Rather, Abbigail Herzog, Brandon Allen, Gabriella Agcoaili, Ciara Parenzin, David Ernst, Mallory Kim, Jarrod Varnell, Taryn Tibbs, Matthew Murphy, Sushant Potdar, Yi Ting Ter, and Kiana Kasmaii. We thank Jessica Proctor for assistance with data curation. Neelandra Joshi, Sarah DuRant, and Jeffrey Lewis provided thoughtful feedback on early versions of these analyses. We thank the Botanical Gardens of the Ozarks for allowing us to gather data on the premises, especially during lockdown. Thank you to the City of Fayetteville for the use of Woolsey Prairie. Special thanks to Pooja Panwar for help locating prairie field sites and for Joe Woolbright at Ozark Ecological Restorations and Arkansas Natural Heritage Commission for permission to survey at Stump and Chesney respectively. This work was funded by an Arkansas Game and Fish Commission Conservation Scholarship to GEH, a Distinguished Doctoral Fellowship through the University of Arkansas to GEH, a University of Arkansas Honors College Research Grant to AEA and ELW, and the University of Arkansas.

## Data Accessibility Statement

Code used to run analyses reported in this article is available on github at XXXXXXX. Data are available on Dryad XXXXXX

## Supplemental Material

All other data presented within this manuscript are available above and as Supplemental Materials.

