## Supplemental Materials for "One size does not fit all: Family specific differences in seasonal patterns of abundance and behavior in butterfly communities"

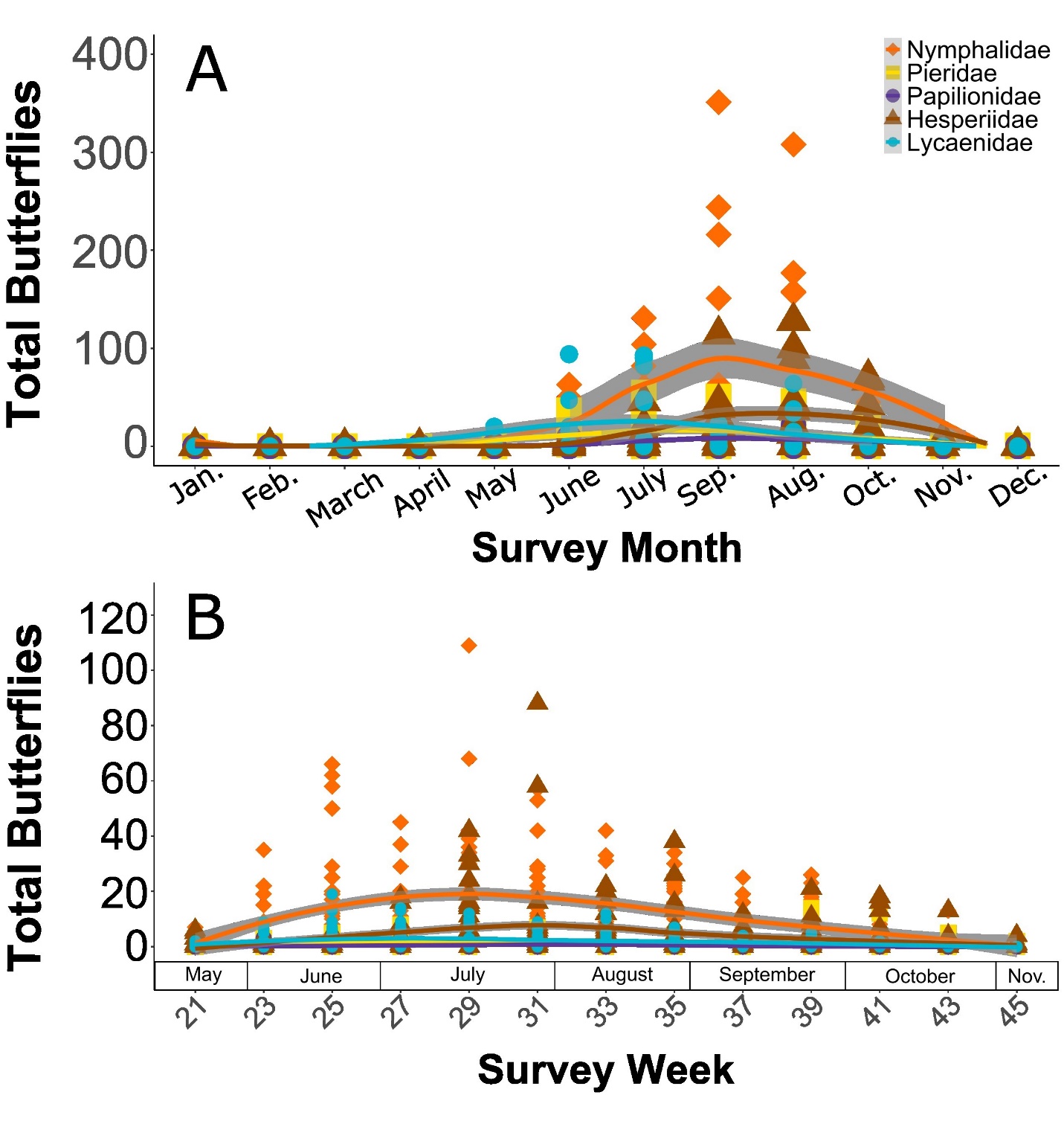

**Supplemental Figure 1**: **Butterfly abundance throughout the year.** Butterfly abundance peaked in August and September in monthly surveys (A) and July in biweekly surveys (B). Monthly surveys were conducted all year round; only the months when butterflies were most abundant are shown.

**Supplemental Table 1**: Correlation matrix of abiotic variables from monthly surveys

|  | Year | Weekly Precipitation | Total Irradiance | Relative UV | Temperature | Month | Site |
| --- | --- | --- | --- | --- | --- | --- | --- |
| Year | 1 | -0.024 | 0.044 | -0.503 | -0.065 | -0.179 | 0.011 |
| Weekly Precipitation | - | 1 | -0.028 | 0.101 | 0.082 | -0.269 | 0.055 |
| Total Irradiance | - | - | 1 | -0.419 | 0.447 | 0.028 | 0.585 |
| Relative UV | - | - | - | 1 | -0.303 | -0.110 | -0.111 |
| Temperature | - | - | - | - | 1 | 0.325 | 0.065 |
| Month | - | - | - | - | - | 1 | 0.021 |

**Supplemental Table 2**: Correlation matrix of abiotic variables from biweekly surveys

|  | Year | Week | Temperature | Weekly Precipitation | Total Irradiance | Relative UV | Site |
| --- | --- | --- | --- | --- | --- | --- | --- |
| Year | 1 | 0.082 | 0.037 | -0.088 | -0.029 | -0.367 | 0.013 |
| Week | - | 1 | -0.466 | 0.013 | -0.303 | -0.580 | -0.043 |
| Temperature | - | - | 1 | -0.217 | 0.464 | 0.118 | 0.015 |
| Weekly Precipitation | - | - | - | 1 | -0.152 | 0.223 | 0.001 |
| Total Irradiance | - | - | - | - | 1 | -0.274 | -0.053 |
| Relative UV | - | - | - | - | - | 1 | 0.011 |
